# Transcriptomics reveals how minocycline-colistin synergy overcomes antibiotic resistance in multidrug-resistant *Klebsiella pneumoniae*

**DOI:** 10.1101/2021.09.21.461326

**Authors:** Thea Brennan-Krohn, Alexandra Grote, Shade Rodriguez, James E. Kirby, Ashlee M. Earl

## Abstract

Multidrug resistant gram-negative bacteria are a rapidly growing public health threat, and the development of novel antimicrobials has failed to keep pace with their emergence. Synergistic combinations of individually ineffective drugs present a potential solution, yet little is understood about the mechanisms of most such combinations. Here, we show that the combination of colistin (polymyxin E) and minocycline has a high rate of synergy against colistin-resistant and minocycline-intermediate or -resistant strains of *Klebsiella pneumoniae*. Furthermore, using RNA-Seq, we characterized the transcriptional profiles of these strains when treated with the drugs individually and in combination. We found a striking similarity between the transcriptional profiles of bacteria treated with the combination of colistin and minocycline at individually subinhibitory concentrations and those of the same isolates treated with minocycline alone. We observed a similar pattern with the combination of polymyxin B nonapeptide (a polymyxin B analogue that lacks intrinsic antimicrobial activity) and minocycline. We also found that genes involved in polymyxin resistance and peptidoglycan biosynthesis showed significant differential gene expression in the different treatment conditions, suggesting possible mechanisms for the antibacterial activity observed in the combination. These findings suggest that the synergistic activity of this combination against bacteria resistant to each drug alone involves sublethal outer membrane disruption by colistin, which permits increased intracellular accumulation of minocycline.

**IMPORTANCE:** Few treatment options exist for multidrug resistant gram-negative bacteria. Synergistic antimicrobial combinations (in which two individually inactive drugs regain activity when used in combination) can overcome resistance. In susceptible bacteria, minocycline acts inside the cell by preventing protein synthesis, while colistin disrupts the outer membrane. Even in bacteria resistant to both drugs, the combination of the two is active. To understand this activity, we studied *Klebsiella pneumoniae* isolates resistant to both drugs. We extracted messenger RNA during treatment with each drug alone and with the combination, then performed RNA sequencing to analyze gene expression levels. Bacteria treated with the combination had an expression profile similar to bacteria treated with minocycline alone. Our findings suggest that synergy results from low-level, subinhibitory membrane disruption by colistin, which permits sufficient accumulation of minocycline in the cell to overcome resistance. Understanding this pathway will facilitate further development of synergistic antimicrobial combination regimens.

## INTRODUCTION

Colistin (also known as polymyxin E), a polypeptide antibiotic that was introduced in 1949, fell out of favor by the 1980s as the result of an unfavorable side effect profile notable for nephrotoxicity and neurotoxicity (1). Rising rates of multidrug resistance among gram-negative bacterial pathogens, including carbapenem-resistant *Enterobacterales* such as *Klebsiella pneumoniae*, subsequently led to a resurgence in the use of colistin due to its broad gram-negative spectrum of activity. Inevitably, resistance to colistin emerged in the setting of increased use, leading to the appearance of pan-drug-resistant isolates (2, 3). Furthermore, colistin’s clinical efficacy is hampered, even in treatment of susceptible isolates, by a low therapeutic index (4). Therefore, there is an urgent need to identify means of restoring the activity of colistin (and the closely related drug polymyxin B) against resistant bacteria and/or reducing the toxicity of the drug while retaining antimicrobial activity.

Structurally, colistin consists of two moieties: a cyclic polypeptide “head” and a lipophilic fatty acid tail [Figure 1]. The drug’s antibacterial activity results from a sequence of events that begins with binding of the polypeptide head to lipopolysaccharide (LPS) on the outer membrane of a gram-negative bacterial cell, leading to outer membrane permeabilization. Colistin exposure ultimately results in disruption of the cytoplasmic membrane, leading to lysis and cell death. The exact mechanisms of these steps remain incompletely understood, but the process is believed to involve self-promoted uptake of the drug, possibly through insertion of the fatty acid tail into the hydrophobic interior of the outer membrane, as well as direct activity at the cytoplasmic membrane itself [Figure 2] (5, 6). Colistin resistance results from modifications to the lipid A component of LPS that increase the charge of LPS, thereby impeding the molecule’s electrostatic interaction with colistin (7). We have previously noted, however, that colistin exhibits robust synergistic activity (defined as a greater-than-additive effect when two drugs are used in combination (8)) against colistin-resistant strains when combined with antibiotics to which the strains are also resistant, including antibiotics that have no intrinsic gram-negative activity (9).

**Figure 1.**
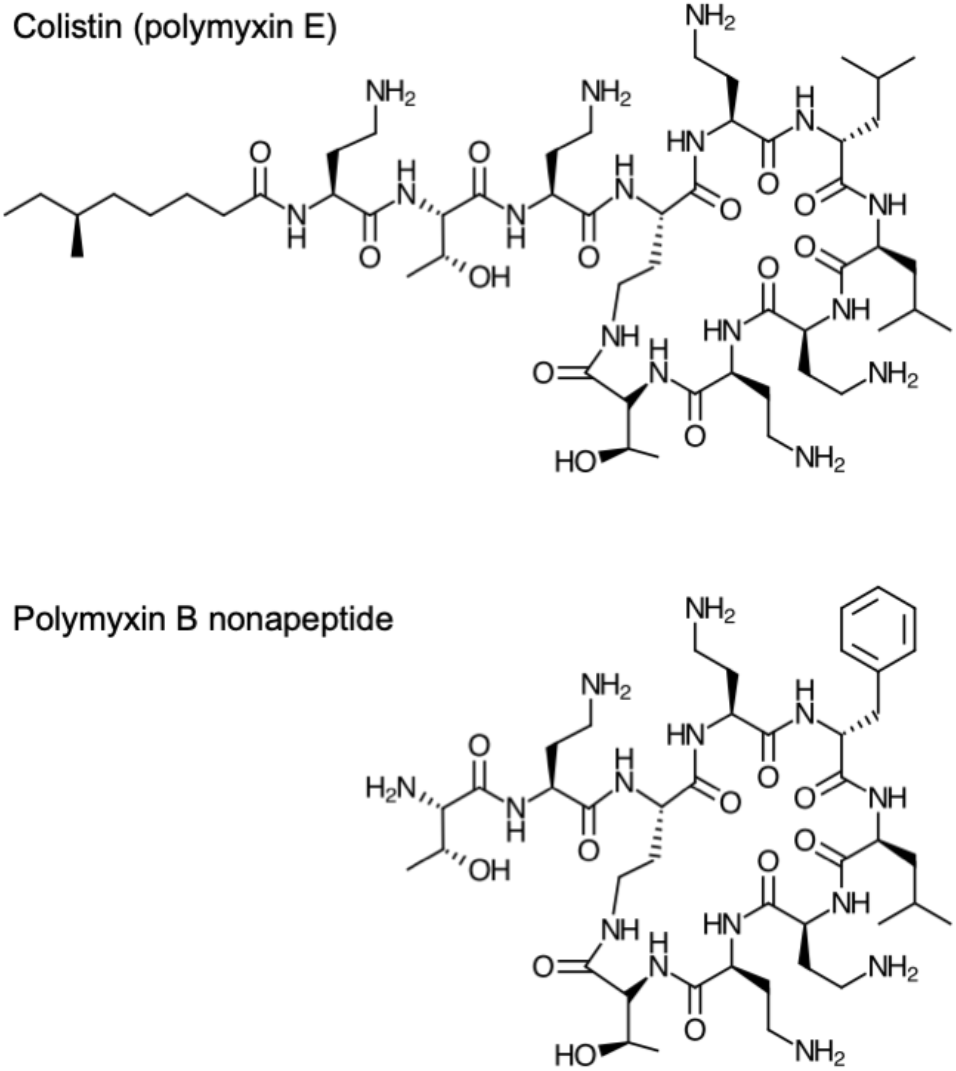
Chemical structure of colistin (polymyxin E) (top), and polymyxin B nonapeptide (PMBN) (bottom). Note the absence of the fatty acid tail in PMBN.

**Figure 2.**
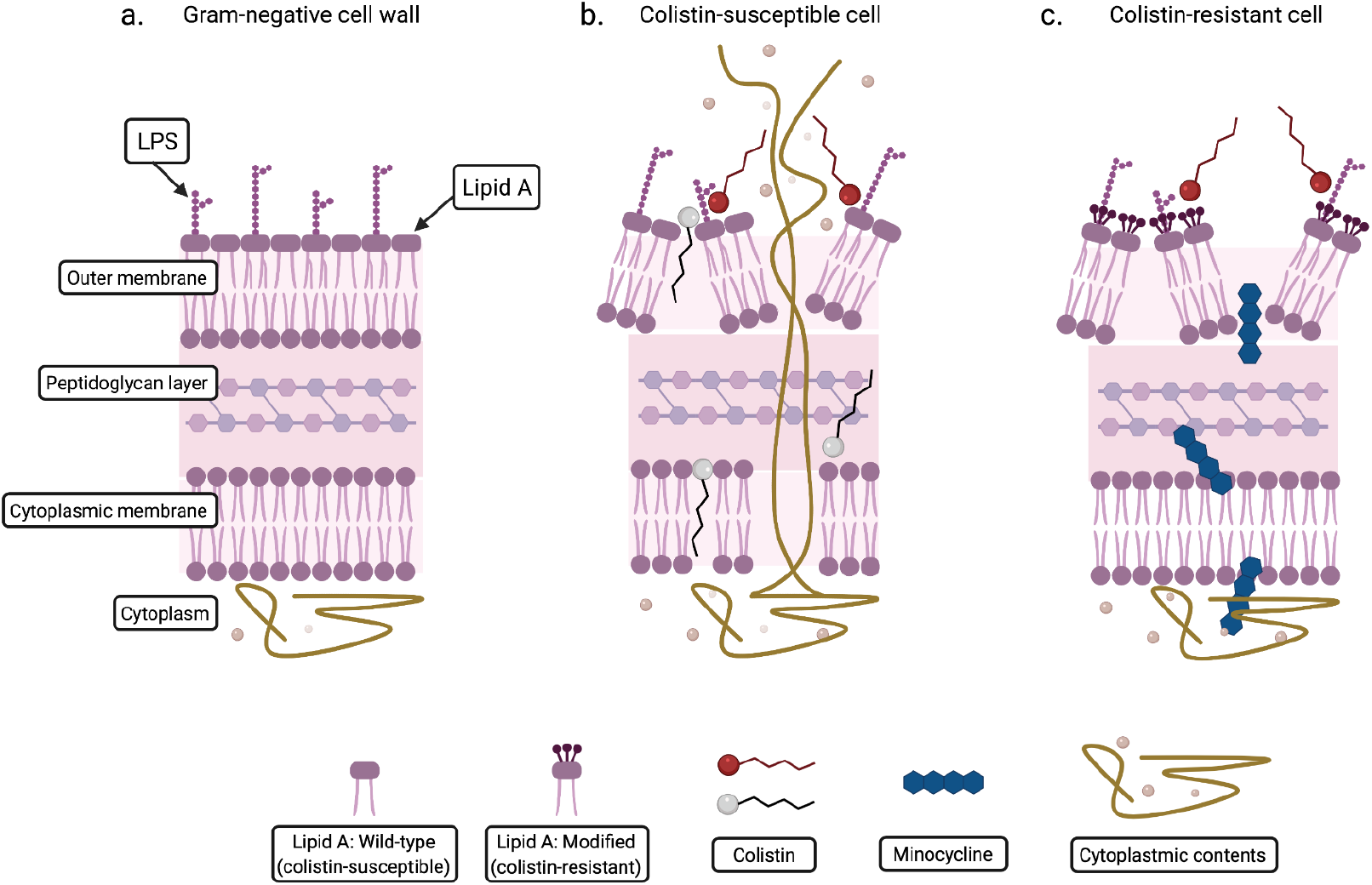
Diagram illustrating the mechanism of action of colistin. A section of gram-negative cell wall is shown (a). When a colistin-susceptible cell is exposed to colistin, permeabilization of the outer membrane, disruption of the cytoplasmic membrane, and cell lysis occur. Red colistin molecules illustrate binding to the lipid A component of LPS. Gray colistin molecules indicate additional proposed activities of colistin, including insertion of fatty acid tail into the membrane interior and activity at the cytoplasmic membrane (b). When a colistin-resistant cell is exposed to colistin, permeabilization of the outer membrane still occurs, allowing increased entry of minocycline. However, disruption of the cytoplasmic membrane and cell lysis do not occur (c).

One of the most consistently synergistic combinations we identified was colistin combined with the tetracycline derivative minocycline (9). This combination is active against the pan-drug resistant “Nevada” *K. pneumoniae* isolate (10) and other isolates that are resistant to both colistin and minocycline, demonstrating synergistic activity at low concentrations of both drugs against these strains. Like other tetracycline antibiotics, minocycline inhibits protein synthesis by binding to the 16S rRNA component of the 30S ribosomal subunit and inhibiting delivery of aminoacyl-tRNAs to the A-site, thereby blocking the elongation step (11). Minocycline has relatively broad intrinsic gram-negative activity, but resistance, primarily due to efflux pump activity, is common.

The sensitization of colistin-resistant bacteria to intracellularly-active antibiotics such as minocycline appears analogous to the capacity of polymyxin B nonapeptide (PMBN) to sensitize gram-negative bacteria to antibiotics that act in the intracellular compartment, despite lacking antibacterial activity on its own (12, 13). PMBN is a derivative of polymyxin B that retains the polypeptide “head” component of the antibiotic but lacks the fatty acid tail [Figure 1]; its activity in combinations is understood to be the result of a retained capacity for membrane disorganization (14). However, the precise mechanism by which loss of the fatty acid tail abrogates direct antibacterial activity while retaining permeabilizing capacity remains incompletely understood.

We hypothesized, by analogy, that synergistic activity in colistin-resistant strains is the result of sub-inhibitory outer membrane permeabilization, which allows increased intracellular accumulation of a drug administered in combination with colistin [Figure 2] (9). In the case of minocycline, increased intracellular accumulation in turn would be sufficient to compensate for minocycline resistance mediated by efflux or other mechanisms. Alternatively, however, it is possible that colistin may have a separate, individually non-inhibitory mechanism of action that is potentiated by the addition of a second antibiotic. It is critically important to elucidate the precise mechanisms of synergy in order to understand how synergistic combinations can be optimized and adopted for clinical use.

We therefore sought to address these alternative possibilities by harnessing the power of transcriptomic technologies. RNA-Seq is a tool that allows quantification of gene expression through sequencing of mRNA transcripts (15) and has been employed in a wide variety of organisms, including bacteria (16). Specifically, RNA-Seq can be used to profile and compare transcriptomic effects resulting from different environmental exposures and as such it should offer a powerful way to compare and contrast the effects of antibiotics used alone or in synergistic combinations, thereby informing our understanding of synergistic activity. To date, there has been minimal use of RNA-Seq to study the activity of colistin or other polymyxin compounds on colistin-resistant bacteria (17, 18), and, to our knowledge, there has been no previous investigation of the effect of colistin-containing combinations on gene expression in colistin-resistant *K. pneumoniae*. We used RNA-Seq transcriptomic analysis to study the effect of the combination of colistin and minocycline on *K. pneumoniae* isolates resistant to both drugs, as an understanding of the mechanism of action of this synergistic combination would provide valuable background and support for its potential clinical application.

## RESULTS

### The combination of minocycline and colistin shows high rates of synergy against colistin-resistant and minocycline-intermediate or -resistant strains using a checkerboard array assay

In order to characterize the activity of the combination of minocycline and colistin across a large number of *K. pneumoniae* strains with different genetic backgrounds and resistance profiles, we performed automated inkjet printer-assisted checkerboard array synergy testing (9,19) on 219 de-identified clinical *K. pneumoniae* isolates [Table S_1], almost all of which have been sequenced (20). The combination was synergistic against 74/219 (34%) of strains overall and against 46/78 minocycline-intermediate or -resistant strains (59%) and 23/25 colistin-resistant strains (92%). Among the 12 strains that were intermediate or resistant to minocycline and resistant to colistin, the combination was synergistic against 10 (83%). For the other two strains, we observed both synergistic and antagonistic wells within the checkerboard array, so the combination was classified as having a mixed rather than synergistic effect. (The two strains in which the mixed phenotype occurred (FDA-CDC 0040 and FDA-CDC 0046; Table S_1) have similar underlying mechanisms for resistance to colistin (*IS1* insertion sequence at nucleotide position 77 of the *mgrB* gene, the gene most commonly mutated in colistin-resistant *K. pneumoniae* (21)) and minocycline (presence of the tet(A) efflux pump). Whole genome comparisons revealed that these two strains are likely clones, having. nearly identical chromosomes (differing only by a single nucleotide polymorphism, indels totalling 18 base pairs and, *IS1* copy number) and the same complement of plasmids.

We also looked for a potentially clinically applicable “salvage” effect among the 10 strains that were intermediate or resistant to both drugs and against which the combination was synergistic. We considered a salvage effect to be present when the concentrations of the two drugs in the synergistic combination were not only reduced from their individually effective concentrations to a degree sufficient to classify them mathematically as synergistic, but were also clinically achievable: in 9/10 of the isolates, the concentrations of minocycline and colistin that were inhibitory in combination were each within the range that would be considered susceptible individually, suggesting potential clinical applicability of the combination for highly multidrug-resistant strains.

### The combination of minocycline and colistin is synergistic in time-kill studies against MGH-149 and BIDMC-32

In order to explore the mechanism responsible for the synergistic killing of multidrug-resistant *K. pneumoniae* by minocycline and colistin, we examined the transcriptional responses of two unrelated colistin- and minocycline-resistant strains [MGH-149 and BIDMC-32; Table 1] that showed the strongest synergistic effect (i.e. lowest minimum fractional inhibitory concentration index (FIC_I_)) among our collection. Both strains carry several multidrug efflux pumps known to be associated with tetracycline resistance (11), and BIDMC-32 also contains the tetracycline-specific efflux pump, TetA [Table S_2]. Likely explaining its colistin resistance, MGH-149 lacks the *mgrB* gene, a negative regulator of the PhoP-PhoQ two-component system, which directs resistance-conferring modification of colistin’s target on lipopolysaccharide molecules (21,22). BIDMC-32 lacks known resistance-conferring mutations in *mgrB* or other genes responsible for lipopolysaccharide modification in *K. pneumoniae*, but does carry a G to T transversion mutation 54 bases upstream of the start codon. This mutation, which is 27 bases downstream of the end of one proposed *mgrB* promoter region (22) and between the −35 and −10 sequences of a second proposed promoter region (23), may affect *mgrB* transcription, potentially through interference with proper promoter binding or activity.

**Table 1.**
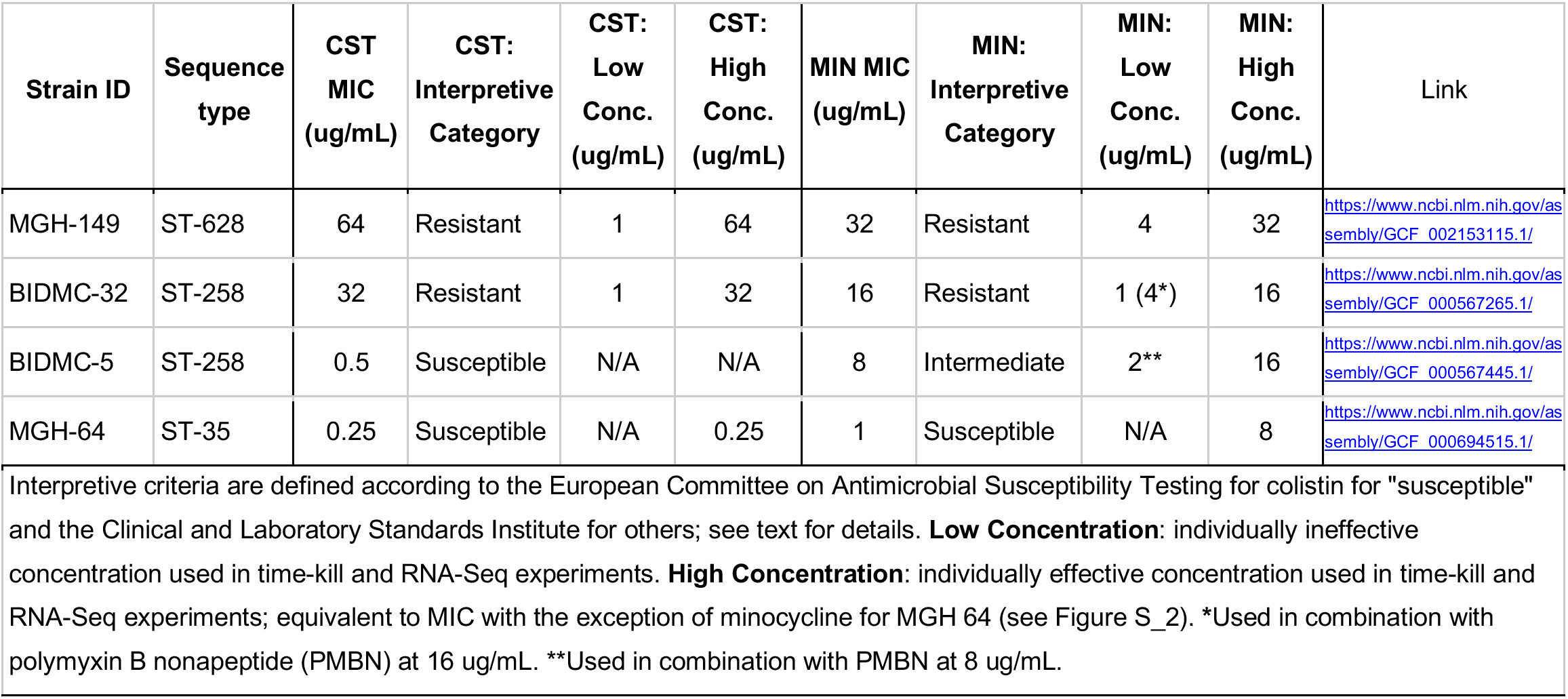
Strains used for transcriptomics

| Strain ID | Sequence type | CST MIC (ug/mL) | CST: Interpretive Category | CST: Low Conc. (ug/mL) | CST: High Conc. (ug/mL) | MIN MIC (ug/mL) | MIN: Interpretive Category | MIN: Low Conc. (ug/mL) | MIN: High Conc. (ug/mL) | Link |
| --- | --- | --- | --- | --- | --- | --- | --- | --- | --- | --- |
| MGH-149 | ST-628 | 64 | Resistant | 1 | 64 | 32 | Resistant | 4 | 32 | https://www.ncbi.nlm.nih.gov/assembly/GCF_002153115.1/ |
| BIDMC-32 | ST-258 | 32 | Resistant | 1 | 32 | 16 | Resistant | 1 (4*) | 16 | https://www.ncbi.nlm.nih.gov/assembly/GCF_000567265.1/ |
| BIDMC-5 | ST-258 | 0.5 | Susceptible | N/A | N/A | 8 | Intermediate | 2** | 16 | https://www.ncbi.nlm.nih.gov/assembly/GCF_000567445.1/ |
| MGH-64 | ST-35 | 0.25 | Susceptible | N/A | 0.25 | 1 | Susceptible | N/A | 8 | https://www.ncbi.nlm.nih.gov/assembly/GCF_000694515.1/ |
| Interpretive criteria are defined according to the European Committee on Antimicrobial Susceptibility Testing for colistin for "susceptible" and the Clinical and Laboratory Standards Institute for others; see text for details. **Low Concentration:** individually ineffective concentration used in time-kill and RNA-Seq experiments. **High Concentration:** individually effective concentration used in time-kill and RNA-Seq experiments; equivalent to MIC with the exception of minocycline for MGH 64 (see Figure S_2). *Used in combination with polymyxin B nonapeptide (PMBN) at 16 ug/mL. **Used in combination with PMBN at 8 ug/mL. |  |  |  |  |  |  |  |  |  |  |

For both strains, we generated synergy time-kill curves in parallel with growth of cells for RNA extraction to evaluate the phenotypic effect of the drugs on bacterial growth over time. Bacteria were treated with colistin and minocycline alone at an individually ineffective (“low”) concentration and with each drug alone at an effective (“high”) concentration, as well as with the two drugs in combination at low concentrations. Concentrations were selected based on results of checkerboard array susceptibility and synergy testing. An untreated control was included in each experiment. At 24 hours, which is the standardized time point for evaluation of bactericidal and synergistic activity in time-kill studies, the combination of minocycline plus colistin was synergistic against both strains (i.e., resulted in a ≥2 log_10_ reduction in CFU/mL at 24 hours compared to the most active agent alone (24)), and was also bactericidal (i.e., resulted in a reduction of ≥3 log_10_ CFU/ml at 24 hours compared to the starting inoculum (25)) against BIDMC-32, but not against MGH-149 [Figure 3].

**Figure 3:**
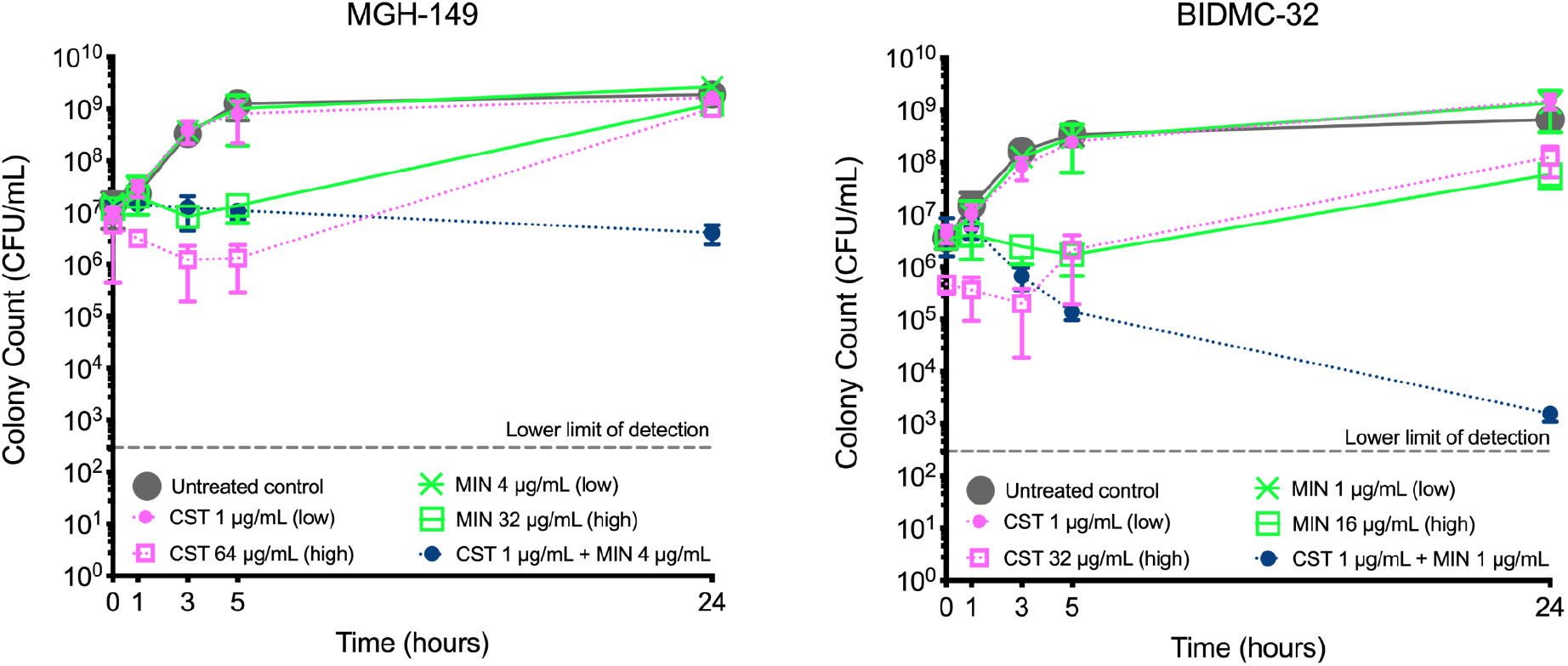
Time-kill curves of MGH-149 and BIDMC-32 treated with minocycline (MIN) and colistin (CST). Results represent the mean and standard deviation of 3 biological replicates. At 2 hours, DNA was extracted from a culture set up under identical conditions in parallel with each replicate. “Low” and “high” refer to the low and high drug concentrations referenced in the text.

In order to understand whether the combination of minocycline with PMBN, a polymyxin drug with outer membrane permeabilizing activity but no direct antibacterial activity, would show similar activity to that of colistin plus minocycline, we tested this combination against BIDMC-32 and against BIDMC-5, a strain that is classified as intermediate to minocycline and susceptible to colistin. The combination was synergistic, although not bactericidal, against both strains in time-kill studies [Figure S_1]. Because we considered that the effects of minocycline and colistin against a resistant strain might be different than their effects against a susceptible isolate, we also tested each drug alone at an individually effective concentration against MGH-64, a representative clinical *K. pneumoniae* isolate that is susceptible to both of these drugs. [Figure S_2]. In several instances, bacteria demonstrated regrowth at 24 hours when treated with one or both drugs at the concentration identified as the minimal inhibitory concentration (MIC) in initial susceptibility and checkerboard array synergy testing, which was performed using the digital dispensing method. The digital dispensing method is an adaptation of broth microdilution testing, and as MICs in the large volumes used in time-kill studies are often higher than MICs determined by broth microdilution testing (26), the observation of regrowth at 24 hours is not unexpected.

### The transcriptomic profile of the synergistic combination is highly similar to that of minocycline alone

We used RNA-Seq to profile the transcriptomes of colistin treatment alone, minocycline treatment alone, and the synergistic combination of colistin and minocycline. From each sample, an average of 11 million reads were generated, 79% of which, on average, aligned to a single reference genome, NC_009648.1/ATCC 700721/MGH 78578, chosen for its similarity to MGH-149 and BIDMC-32 and allowing for cross-strain transcriptomic comparisons (20, 27, 28). Read counts were assigned to genes and other genomic features to generate the transcriptomic profile for each treatment. We performed unsupervised learning using principal component analysis (PCA) on the transcriptomic profiles of each treatment in order to determine how the transcriptional responses to different drugs and drug combinations compared to one another. We found that, in both MGH-149 and BIDMC-32, the transcriptomic responses to the synergistic combination of minocycline and colistin at low concentrations clustered tightly with that of minocycline at high concentration [Figure 4]. Reflective of the minimal effect individual drugs at low concentrations had on the growth of these resistant strains, each strain’s transcriptomic response to exposure to low concentrations of either colistin or minocycline tended to cluster more closely with the untreated controls, while responses to colistin treatment at the high concentration tended to separate from both of these clusters by both principal components [Figure 4]. Principle components one and two accounted for 77% and 16% of the variance in MGH-149, and 66% and 21% of the variance in BIDMC-32, respectively. In both strains, high-dose minocycline treatment alone clusters tightly with the synergistic combination and away from the rest of the treatments along the first principle component.

**Figure 4:**
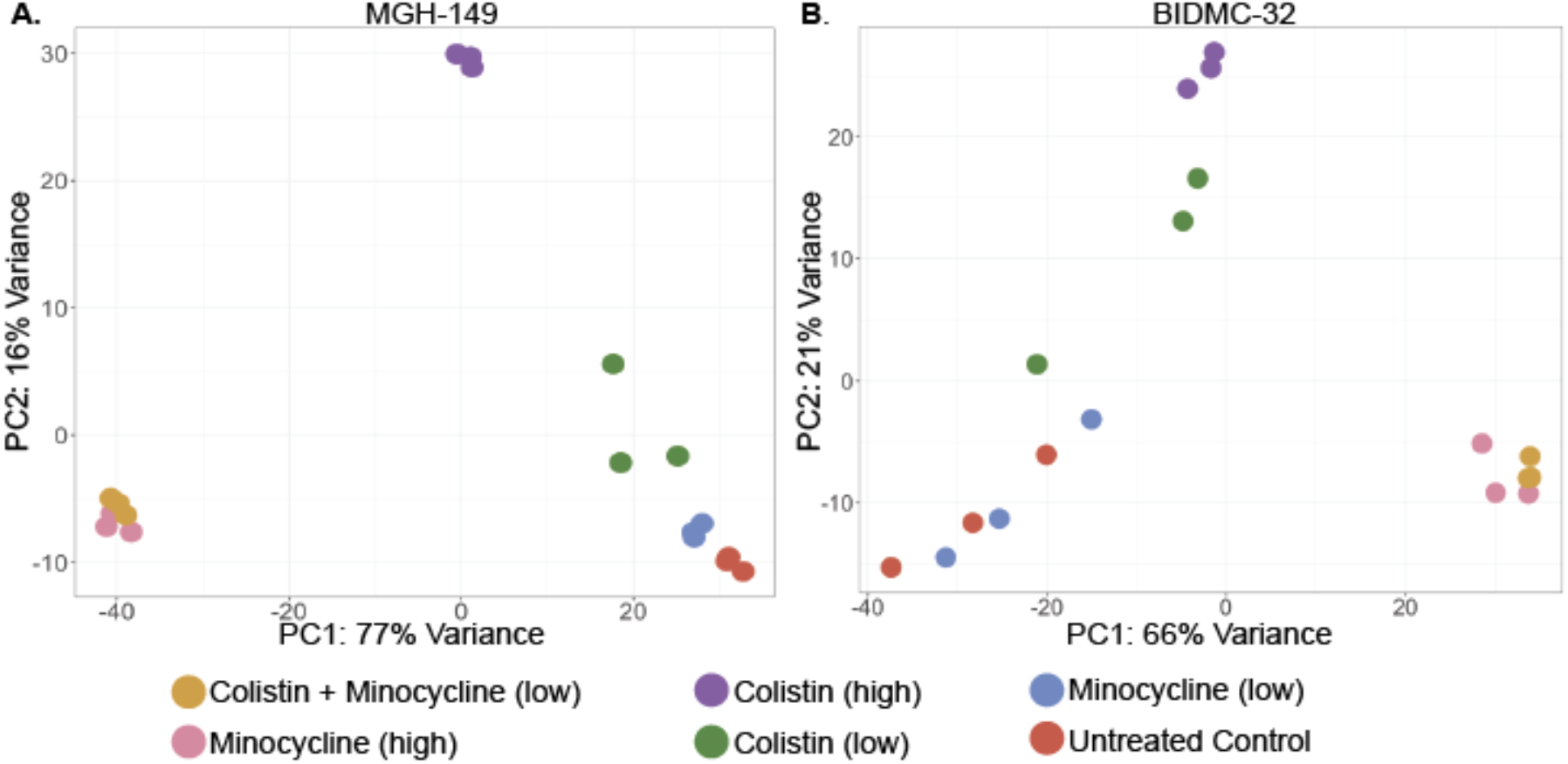
Principal component analysis of the gene expression profiles for antibiotic-treated and the untreated control *K. pneumoniae.* A) PCA of MGH-149 and B) PCA of BIDMC-32. “Low” and “high” refer to the low and high drug concentrations referenced in the text.

Hierarchical clustering of the treatment groups based on gene expression again showed close clustering of the synergistic combination and minocycline at high concentration, as well as shared patterns of co-expressed genes [Figure S_3A (MGH-149) and Figure S_3B (BIDMC-32)].

We next determined the differentially expressed genes for all antibiotic-treated samples compared to the untreated control using DEseq2 (29) [Tables S_3, S_4]. In both strains, the synergistic combination yielded the highest number of differentially expressed genes, followed by minocycline treatment at high concentration. For both strains, the majority of upregulated genes (87% MGH-149; 88% BIDMC-32) and downregulated genes (86% MGH-149; 87% BIDMC-32) in the synergistic combination were also upregulated and downregulated, respectively, at high minocycline concentration [Figure 5, Figure 6]. There were fewer overlaps between differentially expressed genes at high colistin concentration and the synergistic combination: less than half (46% MGH-149; 13% BIDMC-32) of upregulated genes and less than a quarter (24% MGH-149; 18% BIDMC-32) of downregulated genes in the synergistic combination were also upregulated and downregulated, respectively, at high colistin concentration [Figure 5, Figure 6]. Similar sets of GO terms were also enriched in the differentially expressed genes of both strains following treatment with minocycline and the synergistic combination, consistent with the high degree of gene expression overlap [Figures S_4A-D].

**Figure 5:**
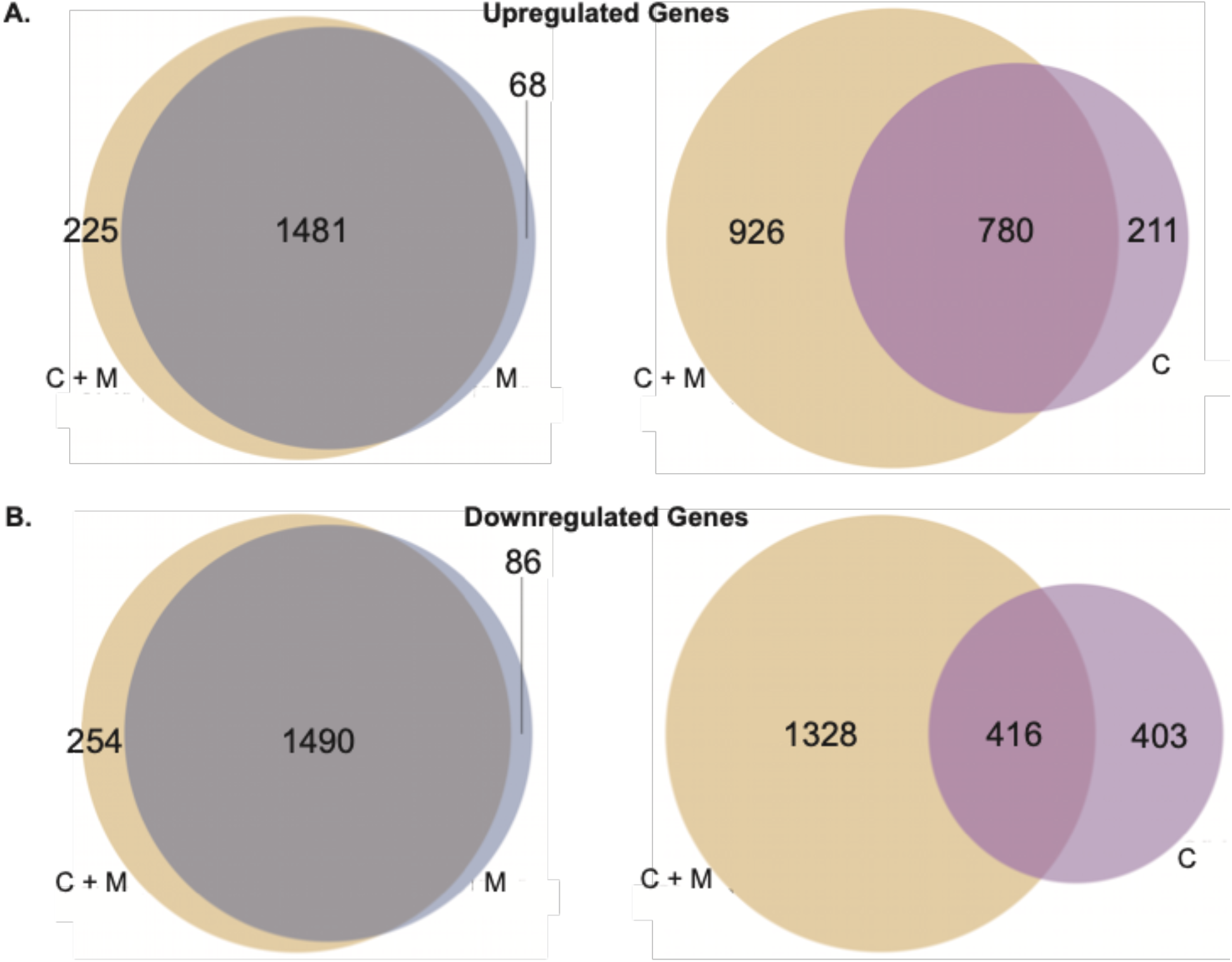
Shared upregulated and downregulated genes between colistin (high) treatment, minocycline (high) treatment and the combination of colistin and minocycline in MGH-149. (A) Upregulated and (B) downregulated differentially expressed genes in each treatment group in *K. pneumoniae* MGH-149 after treatment with colistin (purple), minocycline (Blue), and the combination of colistin and minocycline (Yellow). All comparisons are to the untreated control.

**Figure 6:**
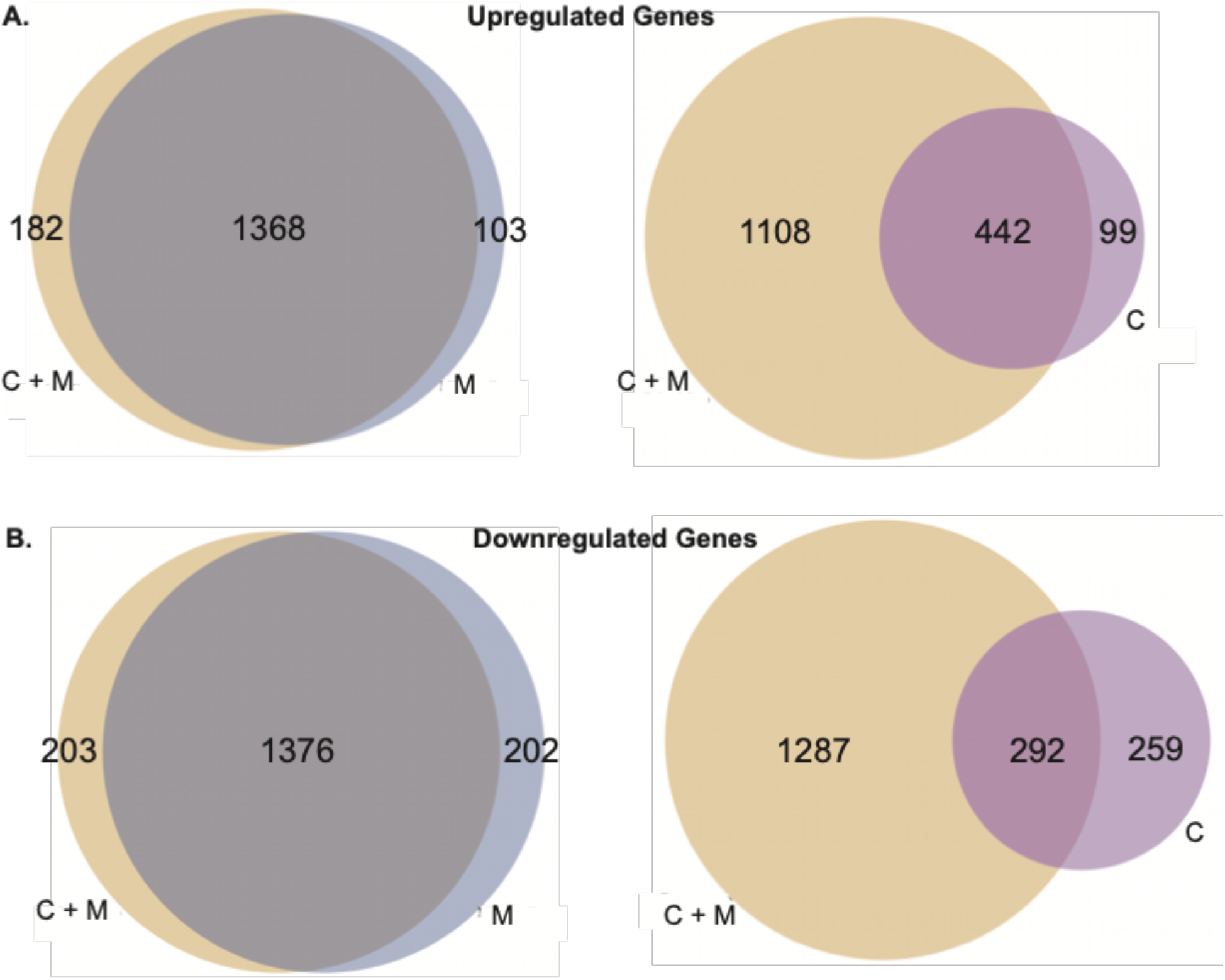
Shared upregulated and downregulated genes between colistin (high) treatment, minocycline (high) treatment and the combination of colistin and minocycline in BIDMC-32. (A) Upregulated and (B) downregulated differentially expressed genes in each treatment group in *K. pneumoniae* BIDMC-32 after treatment with colistin (purple), minocycline (Blue), and the combination of colistin and minocycline (Yellow). All comparisons are to the untreated control.

GO-term enrichment of the downregulated genes in both the synergistic combination and high concentration minocycline treatment was characterized by terms related to metabolic and biosynthetic processes. In contrast, GO-term enrichment of downregulated genes in high concentration colistin treatment was dominated by ion transport functions, including transmembrane transport, as well as membrane-related terms [Figures S_4B and S_4D]. These different enrichment patterns may reflect the inhibition of protein synthesis by minocycline on the one hand, and the impairment of outer membrane integrity by colistin on the other. GO-term enrichment of upregulated genes demonstrated enrichment of genes involved in biosynthesis and metabolism for all treatment conditions, suggesting that upregulated genes, at least in the conditions examined here, may be less reflective of specific antibiotic effects, and instead reflect a more generalized stress response [Figures S_4A and Figure S_4C].

### Genes involved in polymyxin resistance show significant differential gene expression

Most colistin resistance in *K. pneumoniae* occurs via the addition of moieties including L-ara4N to lipid A. This modification decreases the net negative charge of lipid A, reducing binding of polymyxin antibiotics such as colistin (7). Treatment of polymyxin-susceptible *K. pneumoniae* with polymyxin B, a drug closely related to colistin, has previously been observed to increase expression of some genes in the *arnBCADTEF* operon (also known as *pmrHFIJKLM* or *pbgPE*) (18, 30, 31), which encodes enzymes responsible for this lipid A modification [Figure 7A]. We investigated the effects of colistin and the colistin-minocycline combination on expression of these genes in colistin-resistant strains. When colistin-resistant MGH-149 and BIDMC-32 were treated with high concentration colistin, genes in the *arnBCADTEF* operon pathway were highly upregulated, whereas upon treatment with the synergistic combination, these genes were significantly downregulated, as they also were upon treatment with high concentration minocycline alone [Figure 7B]. By contrast, genes in the *arnBCADTEF* operon pathway were not upregulated during exposure to colistin in a colistin-susceptible isolate [Figure S_5]. Taken together, these findings suggest that the subinhibitory permeabilizing activity of low-concentration colistin, as part of combination treatment with minocycline, does not induce pathways promoting colistin resistance in contrast to treatment with high concentration colistin alone in colistin-resistant bacteria. We also examined expression of *pmrD,* which encodes a protein that responds to PhoP signaling by stimulating the two-component system PmrAB (31), which in turn upregulates the *arnBCADTEF* operon. On treatment with high concentrations of colistin, we did not observe increased expression of *pmrD*, suggesting direct activation of *arnBCADTEF* by phoP (Fig. 7), as has been described in *K. pneumoniae* (23, 32). Conversely, *pmrD* was upregulated to varying degrees during treatment with minocycline and with the colistin plus minocycline combination, but without associated upregulation of *arnBCADTEF*.

**Figure 7:**
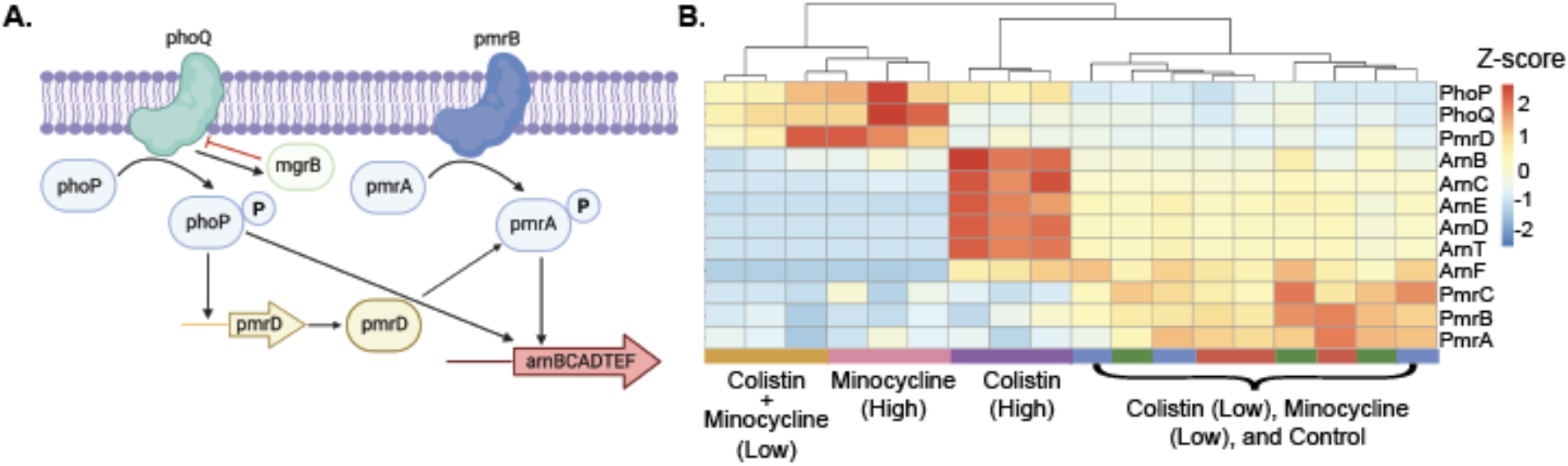
Differential gene expression in the *arnBCADTEF* operon and other genes involved in polymyxin resistance. A) Schematic of the PhoPQ and PmrAB two-component systems and how they control expression of the *arn* operon. Activation of *arnBCADTEF* mediated through PmrD relay of PmrAB signaling, a well as direct activation by PhoP have been described (31, 32) B) Heatmap of gene expression of the pathway in MGH-149 using normalized CPMs showing clustering of biological replicates. Expression was scaled using Z-score normalization prior to clustering, with red representing high expression and blue representing low expression. “(Low)” and “(High)” refer to the low and high drug concentrations referenced in the text.

### The synergistic combination downregulates cell envelope biogenesis

Neither colistin nor minocycline are generally understood to act on synthesis of peptidoglycan in the cell envelope. However, the combination of polymyxin B with chloramphenicol (an antibiotic which, like minocycline, targets protein synthesis in the ribosome) was recently shown to downregulate genes involved in peptidoglycan synthesis (18). To see if the combination of minocycline and colistin would have a similar effect, we evaluated the expression of genes in these pathways. The synergistic combination and high concentration minocycline alone, but not high concentration colistin alone, significantly downregulated the *murB*, *murC*, *murD*, *murE*, *murl*, and *murJ* genes involved in peptidoglycan synthesis. [Figure S_6]. This effect suggests that inhibition of cell envelope biosynthesis may play an unexpected role in the activity of minocycline and the combination, perhaps as a general result of inhibition of protein synthesis.

### The combination of PMBN and minocycline also has a transcriptomic profile similar to minocycline alone

We observed that PMBN was also synergistic with minocycline against both colistin-resistant and colistin-susceptible isolates. PMBN has theoretical therapeutic advantages over colistin in terms of potential reduced toxicity, and PMBN analogs have been explored for clinical use as antibiotic potentiators (33, 34). We were therefore interested in assessing how similar the transcriptional response to the combination of PMBN and minocycline was to the combination of colistin and minocycline. PCA showed clustering of the PMBN and minocycline combination with that of colistin and minocycline, as well as with high concentration minocycline alone, suggesting similar mechanisms for all three treatment conditions [Figure 8].

**Figure 8:**
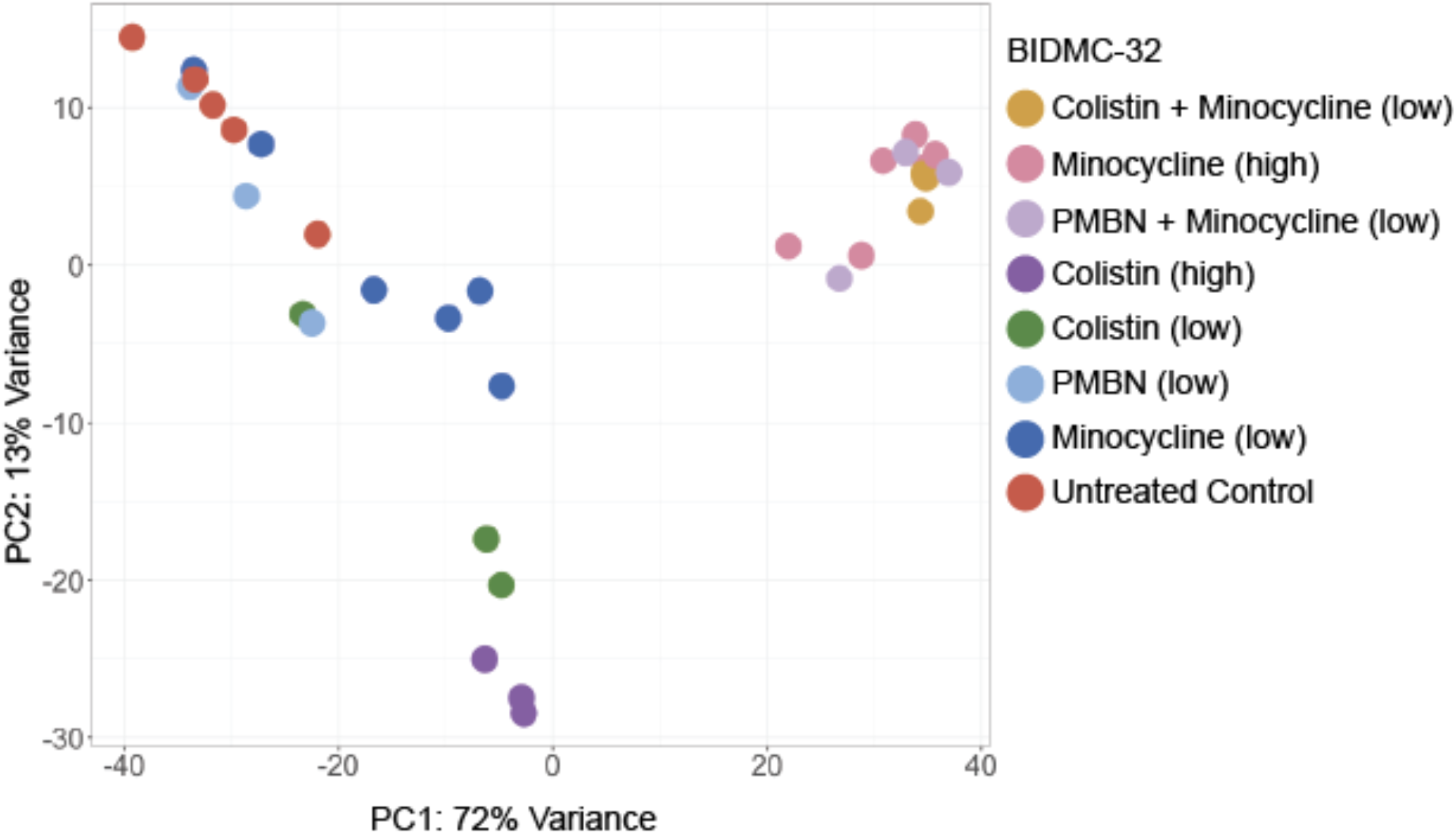
Principal component analysis of the gene expression profiles for antibiotic-treated and untreated control *K. pneumoniae* strain BIDMC-32. “Low” and “high” refer to the low and high drug concentrations referenced in the text. PMBN does not inhibit bacterial growth at any concentration tested (it does not have an MIC) and therefore was tested both alone and in combination with minocycline at its FIC (“low”).

## DISCUSSION

Polymyxins, including colistin, have long been recognized as outer membrane-disrupting agents that facilitate uptake of hydrophobic antibiotics that are normally excluded from the interior of gram-negative cells as a result of their inability to cross the outer membrane (35). In colistin-susceptible bacteria, this outer membrane permeabilization effect is combined with cytoplasmic membrane disruption, resulting in cell lysis and death. Recently, studies describing synergistic activity of colistin-containing antibiotic combinations against colistin-resistant bacteria have provided evidence that colistin retains outer membrane permeabilizing activity against colistin-resistant bacteria, despite its absence of lytic activity or growth inhibition in these strains (9, 36). An analogous phenomenon of isolated permeabilization occurs when bacteria are treated with PMBN, which retains the outer membrane permeabilizing effect but not the cytoplasmic membrane disrupting effect of its parent compound (37). In the results presented here, we observed a striking similarity between the transcriptomic profiles of colistin- and minocycline-resistant *K. pneumoniae* isolates treated with a combination of colistin and minocycline at individually subinhibitory concentrations and the profiles of those same isolates treated with minocycline alone, and with PMBN plus minocycline. These findings support the hypothesis that the synergistic activity of the colistin plus minocycline combination involves sublethal cell envelope disruption by colistin, which permits increased intracellular accumulation of minocycline. Tetracycline antibiotics can enter gram-negative cells either through porin channels or by diffusion through the outer membrane lipid bilayer (38), with outer membrane diffusion playing a larger role for hydrophobic tetracycline derivatives like minocycline (39). It is thus logical that increased permeability of the outer membrane would have a marked effect on minocycline entry into the cell and that an increased rate of entry could overcome drug efflux (40), which is the primary mechanism of minocycline resistance in the strains evaluated here.

While the explanation for the decoupling of outer membrane permeabilization and cell lysis in colistin-resistant bacteria is not fully understood (41), it may involve the presence of lipid A molecules at the cytoplasmic membrane that undergo resistance-conferring modifications at a higher rate relative to the outer membrane, leading to decreased vulnerability of the cytoplasmic membrane to the effect of colistin (42). Loss of the fatty acid tail in PMBN results in a phenotypically similar effect to the impaired polypeptide “head” binding of colistin in colistin-resistant bacteria, yet the exact explanation for the loss of lytic activity in PMBN also remains incompletely elucidated. It has been proposed that self-promoted uptake of colistin across the outer membrane involves insertion of the fatty acid tail into the outer membrane (5). Lacking this tail, PMBN would be unable to reach the cytoplasmic membrane, while its permeabilizing effect at the outer membrane would remain unaffected. Consistent with this hypothesis, PMBN has been shown to cause leakage of periplasmic but not cytoplasmic proteins (43). One report describes loss of K+, uracil, and amino acids across the cytoplasmic membrane during treatment with PMBN, but the rate of loss was less than during treatment with polymyxin B and was not associated with bacterial killing (6). Interestingly, PMBN treatment causes less morphological outer membrane damage than polymyxin B, suggesting that there may in fact also be differences between PMBN and intact polymyxin compounds in terms of outer membrane disruption (43). We observed bacteriostatic synergistic activity against both colistin-resistant and colistin-susceptible strains when PMBN was combined with minocycline [Figure S_1] (in contrast to bactericidal activity of colistin and minocycline against one of the same strains), and noted that treatment of the colistin-resistant strain BIDMC-32 with minocycline and PMBN resulted in a transcriptomic profile that clustered tightly with that of the combination of minocycline and colistin [Figure 8]. These observations do not fully disentangle the relative roles of the polypeptide “head” and fatty acid tail in the activity of polymyxins and polymyxin derivatives at the outer membrane and cytoplasmic membrane. However, they do provide further evidence of the highly similar inhibitory process effected by PMBN and, in colistin-resistant bacteria, by colistin, when combined with intracellularly active antibiotics.

We observed that genes in the *arnBCADTEF* operon were upregulated during treatment with colistin alone and downregulated during treatment with minocycline alone or the combination of colistin and minocycline. A similar pattern was observed by Abdul Rahim and colleagues in a polymyxin B-susceptible *K. pneumoniae* isolate treated with polymyxin B, chloramphenicol (a protein synthesis inhibitor antibiotic), and the combination (18). The *arnBCADTEF* operon mediates production of 4-amino-4-deoxy-L-arabinose (L-Ara4N) and addition of this molecule to lipid A, which causes a reduction in the negative charge of lipid A and a resultant inability of colistin to bind to this target (7). Modification of lipid A by L-Ara4N is the most common mechanism of colistin resistance in *K. pneumoniae* (44) and the expected mechanism in strains MGH-149 (as a result of loss of the *mgrB* repressor gene) and BIDMC-32 (possibly through a point mutation located between the promoter and start codon of the *mgrB* gene). The PhoP/PhoQ two-component system, which operates upstream of expression of the *arnBCADTEF* operon, is known to be responsive to antimicrobial peptides (45) and has been shown to be upregulated by polymyxin B exposure in *Pseudomonas aeruginosa* (46), so it is possible that activation of PhoP/PhoQ in response to colistin results in downstream upregulation of the *arnBCADTEF* operon in a feed-forward pattern of colistin-responsive colistin resistance. Signaling from PhoPQ can be transmitted to the *arnBCADTEF* operon by the connector protein PmrD and the PmrAB two-component system (47) (Figure 7), but we did not observe significant upregulation of *pmrD* during colistin treatment. This may reflect upregulation of *arnBCADTEF* through direct signaling from PhoPQ, which has been described in *K. pneumoniae* (32) as well as *Serratia marcescens* (48); indeed, a recent proteomics analysis suggested that direct activation by PhoPQ may be the primary means of *arnBCADTEF* upregulation in *K. pneumoniae* under growth conditions similar to those used in this work. (23) (We note that neither *phoP/phoQ* nor *pmrA/pmrB* were upregulated during colistin treatment, but we would not necessarily expect activation of these two-component systems to result in changes in their gene expression). We did not observe upregulation of the *arnBCADTEF* operon in response to colistin exposure in the colistin-susceptible strain MGH-64 [Figure S_5].

Upregulation of the *arnBCADTEF* operon did not occur during exposure to minocycline alone or to the synergistic combination of minocycline and colistin, suggesting that the subinhibitory action of low concentrations of colistin on colistin-resistant bacteria does not induce its own resistance. This finding, along with the observation that the combination, but not either drug alone, prevented regrowth at 24 hours in the time-kill experiments, suggests that the combination may be more resilient against the emergence of bacterial resistance or tolerance. We did observe upregulation of *pmrD* during exposure to minocycline and the minocycline/colistin combination, but this increase in transcription of the upstream two-component system did not ultimately result in downstream upregulation of the *arnBCADTEF* operon under these treatment conditions. The lack of correlation between *pmrD* and *arnBCADTEF* expression under these conditions might suggest an absence of pmrAB signaling, such that increased levels of *pmrD* are not able to affect *arnBCADTEF* expression via enhancement of the pmrAB two-component system.

The development of novel antimicrobials over the past several decades has failed to keep pace with the emergence of multidrug resistance, particularly among gram-negative bacteria (49). Colistin has traditionally been considered a drug of last resort for treatment of highly resistant gram-negative pathogens, but it is not a useful treatment option for the increasing number of colistin-resistant isolates (50), and its narrow therapeutic window limits its clinical utility in treating even colistin-susceptible strains (51). Synergistic combinations of individually ineffective drugs, as evaluated here, offer potential therapeutic alternatives that do not depend on the development of new antibacterial agents. Indeed, a PMBN-like compound with outer membrane permeabilizing activity has been evaluated as an antibiotic potentiator against multidrug-resistant gram-negative pathogens and has undergone Phase I investigation (34, 52). Although efficacy studies in human subjects will ultimately be required to assess the clinical efficacy of such combinations, *in vitro* investigations such as we present here can inform such studies by helping to elucidate the mechanism of action and indicating plausible dosing regimens.

Transcriptomic studies can provide insight into a wide range of cellular processes by comparing large, systematically acquired datasets, but they also possess certain intrinsic limitations. Although we can identify genes and gene pathways that are up- or down-regulated, we cannot, for example, quantify translation or protein levels. However, identifying significantly differentially expressed genes can help shed light on partially understood processes and allows us to prioritize pathways for further evaluation using complementary techniques.

We present here a novel use of this promising technology to further our understanding of antimicrobial combination therapy for treatment of some of the most highly drug-resistant pathogens identified to date. Many antibiotics still in common use, including tetracyclines and polymyxins, as well as β-lactam drugs (e.g. penicillin), were discovered and entered clinical use long before the scientific tools necessary to elucidate their mechanisms of action had been developed. As a result, much still remains unknown about the spectrum of effects that each of these compounds exerts on bacteria, and RNA-Seq analysis, as we have demonstrated here, can provide a valuable tool for enhancing our understanding of these drugs.

## MATERIALS AND METHODS

### Bacterial strains

*Klebsiella pneumoniae* strains were obtained from the carbapenem-resistant *Enterobacterales* genome initiative at the Broad Institute (Cambridge, MA), (213 strains) (20) and the U.S. Food and Drug Administration (FDA)-CDC Antimicrobial Resistance Isolate Bank (Atlanta, GA; 6 strains). A full list of strains is included in Table S1. *Escherichia coli* ATCC 25922 was obtained from the American Type Culture Collection (Manassas, VA). All strains were colony purified, minimally passaged, and stored at 80°C in tryptic soy broth with 50% glycerol (Sigma-Aldrich, St. Louis, MO) until the time of use.

### Antibiotics

Colistin sulfate was obtained from Santa Cruz Biotechnology (Santa Cruz, CA) and minocycline hydrochloride was obtained from Chem-Impex International (Wood Dale, IL). Antibiotic stock solutions for time-kill studies and for growth of bacteria for RNA extraction were prepared in water; stock solutions used for the digital dispensing method were prepared in water with 0.3% polysorbate 20 (P-20; Sigma-Aldrich, St. Louis, MO) as required by the HP D300 digital dispenser instrument (HP, Inc., Palo Alto, CA) for proper fluid handling. All antibiotic stock solutions were quality control tested with *E. coli* ATCC 25922 by standard broth microdilution technique using the direct colony suspension method (53) or the digital dispensing method (for stocks to be used for checkerboard arrays) prior to use. Antimicrobial solutions were stored as aliquots at 20°C and were discarded after a single use.

### Checkerboard array synergy testing and MIC determination

To create checkerboard arrays, serial 2-fold dilutions of colistin (concentration range 0.016 - 16 μg/mL) and minocycline (concentration range 0.008 - 32 μg/mL) were dispensed in orthogonal titrations by the D300 instrument using the digital dispensing method previously developed by our group (54, 55). Bacterial inocula were prepared by suspending colonies in cation-adjusted Mueller-Hinton broth (CAMHB; BD Diagnostics, Franklin Lakes, NJ) and diluting to a final concentration of 5×10^5^ CFU/mL. Antimicrobial stocks were dispensed in appropriate volumes into empty, flat-bottomed, untreated 384-well polystyrene plates (Greiner Bio-One, Monroe, NC) by the D300. Immediately after addition of antimicrobials, the wells were inoculated with 50 μL of bacterial suspension and the plates were incubated for 16 to 20 h at 35°C in ambient air. After incubation, bacterial growth was quantified by measurement of optical density at 600 nm (OD_600_) using a Tecan Infinite M1000 Pro microplate reader (Tecan, Morrisville, NC). An OD600 reading of 0.07 or greater (concordant with visual growth assessment) was considered indicative of bacterial growth. Minimal inhibitory concentrations (MICs) for each antibiotic were determined from wells in the array containing only that drug. Isolates were considered susceptible, intermediate, or resistant according to CLSI breakpoint tables (56), with the exception of the classification of susceptible for colistin, as CLSI does not have a “susceptible” category for colistin for *Enterobacterales*; isolates with MIC ≤2 μg/mL are classified as “intermediate” to reflect colistin’s limited clinical efficacy. For clarity in the manuscript, we used the current European Committee on Antimicrobial Susceptibility Testing (EUCAST) interpretive criteria for colistin for *Enterobacterales*, which classifies isolates with an MIC ≤2 as susceptible (57). For each concentration of each drug, the well in which growth was inhibited at the lowest concentration of the other drug was identified. For each of these wells, the fractional inhibitory concentration (FIC) for each antibiotic was calculated by dividing the concentration of the antibiotic in that well by the MIC of the antibiotic. The FIC index (FIC_I_) for each well was then determined by summing the FICs of the two drugs. If the MIC of an antibiotic was off scale, the highest concentration tested was assigned an FIC of 0.5 to permit FIC_I_ calculation. When the minimum FIC_I_ in a synergy grid was ≤0.5, the combination was considered synergistic for that bacterial isolate, and when the maximum FIC_I_ was >4.0, the combination was considered antagonistic. When all FIC_I_ values were intermediate, the combination was considered to have no interaction. In cases where a single grid included both synergistic and antagonistic FIC_I_ values, the combination was considered to have a mixed effect.

### Growth conditions for RNA extraction

Time-kill synergy studies and RNA extraction for RNA-Seq were performed using the following *K. pneumoniae* isolates and treatment conditions (Table 1): BIDMC-32, exposed to minocycline and colistin and to minocycline and polymyxin B nonapeptide, separately and in combination, MGH-149, exposed to minocycline and colistin, separately and in combination, BIDMC-5, exposed to minocycline and polymyxin B nonapeptide, separately and in combination, and MGH-64, exposed to minocycline and colistin separately. Antibiotic concentrations in each condition were selected based on results of checkerboard array synergy testing, with final concentrations adjusted as necessary, based on preliminary time-kill experiments, to account for expected differences in MIC and FIC concentrations between the microdilution volumes used in checkerboard array experiments (50 μL per reaction) and in time-kill experiments (15 mL per reaction). Antibiotic stocks were prepared in water and constituted a volume of ≤3.2% of the total volume of the culture; because the carrier volume was small and consisted only of water, blank carrier liquid was not added to untreated controls. For each antibiotic/bacterial strain condition, two identical culture tubes were prepared in parallel from the same starting inoculum, with one used for the time-kill study and one used for RNA extraction. All studies were performed with three biological replicates.

Antibiotic stocks were prepared as described above and diluted in 15 mL of CAMHB in 25-by 150-mm glass round-bottom tubes to the desired starting concentrations. An initial starting inoculum was prepared by adding 250 μL of a 0.5 McFarland suspension of colonies from an overnight plate to 12.5 ml of CAMHB and incubating on a shaker in ambient air at 35°C until the suspension reached log-phase growth (approximately 4 h). The culture was then adjusted to the turbidity of a 1.0 McFarland standard in CAMHB, and 0.75 mL of this suspension was added to each of the tubes containing antimicrobial solutions for a final starting inoculum of ~5×10^6^ CFU/mL. An untreated control and a negative control tube were prepared in parallel with each experiment. Cultures were incubated on a shaker in ambient air at 35°C. For RNA extraction, one set of tubes was removed from the incubator at 2 hours. The contents of each tube was spun at 2000 xg for 20 min at 4°C. The supernatant was removed and the pellet was resuspended in 0.5 mL of TRIzol reagent (Invitrogen, Waltham, MA), incubated at room temperature for 5 minutes, then frozen at −80°C until the time of RNA extraction. For the time-kill study, aliquots from each culture tube from the second set of tubes were removed for colony enumeration at 0, 1, 3.5, and 24h (MGH-64) and at 0, 1, 3, 5, and 24h (other strains). To perform colony counts, a 10-fold dilution series was prepared in 0.9% sodium chloride. A 10 μL drop from each dilution was transferred to a Mueller-Hinton plate (Thermo Fisher, Waltham, MA) and incubated overnight in ambient air at 35°C (58). For countable drops (drops containing 3 to 30 colonies), the cell density of the sample was calculated; if more than one dilution for a given sample was countable, the cell density of the two dilutions was averaged. If no drops were countable, the counts for consecutive drops above and below the countable range were averaged. The lower limit of detection was 300 CFU/mL.

### RNA extraction

Cell pellets resuspended in 0.5 mL Trizol reagent (ThermoFisher Scientific) were transferred to 2mL FastPrep tubes (MP Biomedicals) containing 0.1 mm Zirconia/Silica beads (BioSpec Products) and bead beaten for 90 seconds at 10 m/sec speed using the FastPrep-24 5G (MP Biomedicals). After addition of 200 ul chloroform, each sample tube was mixed thoroughly by inversion, incubated for 3 minutes at room temperature, and spun down 15 minutes at 4°C. The aqueous phase was mixed with an equal volume of 100% ethanol, transferred to a Direct-zol spin plate (Zymo Research), and RNA was extracted according to the Direct-zol protocol (Zymo Research).

### Generation of RNA-Seq data

Illumina cDNA libraries were generated using a modified version of the RNAtag-seq protocol (59, 60). Briefly, 500ng-1 μg of total RNA was fragmented, depleted of genomic DNA, dephosphorylated, and ligated to DNA adapters carrying 5’-AN_8_-3’ barcodes of known sequence with a 5’ phosphate and a 3’ blocking group. Barcoded RNAs were pooled and depleted of rRNA using the RiboZero rRNA depletion kit (Epicentre). Pools of barcoded RNAs were converted to Illumina cDNA libraries in 2 main steps: (i) reverse transcription of the RNA using a primer designed to the constant region of the barcoded adaptor with addition of an adapter to the 3’ end of the cDNA by template switching using SMARTScribe (Clontech) as described [3]; (ii) PCR amplification using primers whose 5’ ends target the constant regions of the 3’ or 5’ adaptors and whose 3’ ends contain the full Illumina P5 or P7 sequences. cDNA libraries were sequenced to generate paired end reads.

### Analysis of RNA-Seq data

Sequencing reads from each sample in a pool were demultiplexed based on their associated barcode sequence using custom scripts (https://github.com/broadinstitute/split_merge_pl). Up to 1 mismatch in the barcode was allowed provided it did not make assignment of the read to a different barcode possible. Barcode sequences were removed from the first read as were terminal G’s from the second read that may have been added by SMARTScribe during template switching. Reads were aligned to NC_009648.1/ATCC 700721/MGH 78578, a multidrug-resistant reference strain originally isolated from the sputum of a patient (20, 27, 28) using BWA (61) and read counts were assigned to genes and other genomic features using custom scripts (https://github.com/broadinstitute/BactRNASeqCount). Read counts were used as input to DESeq2 (Release 3.12) to build a DEseq dataset [30]. Differential gene expression analysis was performed using DESeq2 default settings. In order to check for batch effects, Principal Component Analysis (PCA) was performed on biological replicates after variance-stabilizing transformation to obtain transformed, normalized counts. Differentially expressed genes were determined between each treatment group and the untreated control. Genes were determined as differentially expressed using a threshold of *p* <0.05 and a false-discovery rate (FDR) of 5%, standard settings in DESeq2. Differential gene expression analysis was also performed using EdgeR (Version 3.32.1) (62, 63) obtaining very similar results to DESeq2. EdgeR was used to calculate CPMs, Counts Per Million, for each gene, which was used for the unsupervised clustering and heatmap generation.

## ACKNOWLEDGEMENTS

RNA-Seq libraries were constructed and sequenced at the Broad Institute of MIT and Harvard by the Microbial ‘Omics Core and Genomics Platform, respectively. The Microbial ‘Omics Core also provided guidance on experimental design and conducted preliminary analysis for all RNA-Seq data. The authors would like to thank Abigail Manson, Roby Bhattacharyya, Noam Shoresh, and Alejandro Pironti for their input and suggestions. T.B.-K. was supported by a Eunice Kennedy Shriver National Institute of Child Health and Human Development pediatric infectious diseases research training grant (T32HD055148), a National Institute of Allergy and Infectious Diseases (NIAID) training grant (T32AI007061), a Boston Children’s Hospital Office of Faculty Development Faculty Career Development fellowship, and a NIAID career development award (1K08AI132716). ATG was supported by the Jane Coffin Childs Memorial Fund for Medical Research. This work was also funded in part with Federal funds from the National Institute of Allergy and Infectious Diseases, National Institutes of Health, Department of Health and Human Services, under Grant Number U19AI110818 to the Broad Institute. The HP D300 digital dispenser and TECAN M1000 used in experiments were provided by TECAN (Morrisville, NC). TECAN had no role in study design, data collection/interpretation, manuscript preparation, or decision to publish. Figures 1 and 7A were created with https://BioRender.com.

